# flexiMAP: A regression-based method for discovering differential alternative polyadenylation events in standard RNA-seq data

**DOI:** 10.1101/672766

**Authors:** Krzysztof J. Szkop, David S. Moss, Irene Nobeli

**Affiliations:** Institute of Structural and Molecular Biology, Department of Biological Sciences, Birkbeck, Malet Street, London WC1E 7HX, UK

## Abstract

**Summary:** We present flexiMAP (flexible Modeling of Alternative PolyAdenylation), a new beta-regression-based method implemented in *R*, for discovering differential alternative polyadenylation events in standard RNA-seq data. Importantly, flexiMAP allows modeling of multiple known covariates that often confound the results of RNA-seq data analysis. We show, using simulated data, that flexiMAP is very specific and outperforms in sensitivity existing methods, especially at low fold changes. In addition, the tests on simulated data reveal some hitherto unrecognised caveats of existing methods.

**Availability:** The flexiMAP *R* package is available at: https://github.com/kszkop/flexiMAP

Scripts and data to reproduce the analysis in this paper are available at: https://doi.org/10.5281/zenodo.3238619

**Contact:** Irene Nobeli

## Introduction

Alternative polyadenylation (APA) is the selection of alternative cleavage and polyadenylation sites during transcription of eukaryotic genes, resulting in isoforms with distinct 3’ untranslated regions. APA has been shown to be prevalent in mammalian transcripts and alternative isoforms are linked to different stages of development, cell types and disease status (Elkon *et al.*, 2013; Szkop *et al.*, 2017). APA events can be identified on a genome-wide scale using 3’ end-focused sequencing or, more recently, long-read sequencing, but as these methods are still not widely used and many legacy transcriptome surveys were carried out using standard RNA-seq sequencing instead, it would be useful to have computational methods that can identify differential APA in RNA-seq data. A few such methods exist already (Xia *et al.*, 2014; Grassi *et al.*, 2016; Ha *et al.*, 2018; Ye *et al.*, 2018; Arefeen *et al.*, 2018) but they have caveats (Szkop and Nobeli, 2017). For example, all methods must solve the problem of how to deal with biological replicates; some test the replicates individually, losing the advantage of having replicates in the first place; others, average values from replicates, effectively losing track of the within-group variability. In designing a method for differential APA analysis, we considered the following: a) the reconstruction and quantification of the individual isoforms is both challenging and not strictly necessary for this task; b) the errors in modeling RNA-seq read counts are neither normal nor Poisson-distributed; c) multiple covariates can affect APA.

Inspired by the use of Generalized Linear Models (GLMs) in differential gene expression (Robinson *et al.*, 2010; Love *et al.*, 2014) we present here a regression-based method and associated pipeline (flexible modeling of APA or flexiMAP) that satisfactorily addresses the above requirements. In addition, we show, using simulated data, that the method is both sensitive and specific across a range of fold changes and numbers of samples and that its performance is superior to three alternatives (DaPars (Xia *et al.*, 2014), Roar (Grassi *et al.*, 2016) and APAtrap (Ye *et al.*, 2018)) in most tests we carried out. The method is available as an *R* package from: https://github.com/kszkop/flexiMAP

## Method Description

Our method can be applied to all pairs of polyadenylation sites in a gene, where one site is considered “distal” (i.e. located furthest away from the end of the coding region) and one is “proximal” (Supp. Fig. 1). The proximal site separates the 3 ‘UTR into two regions: the “short” region, starting at the end of the coding region and ending at the proximal site, and the “long” region starting at the proximal site and ending at the end of the transcript (Supp. Fig. 1). Assuming the separation of samples into groups based on the condition of interest, the question we want to answer is: given a total number of reads falling in the 3’ UTR, is the proportion of reads falling in the long region dependent on the sample group membership?

**Figure 1.**
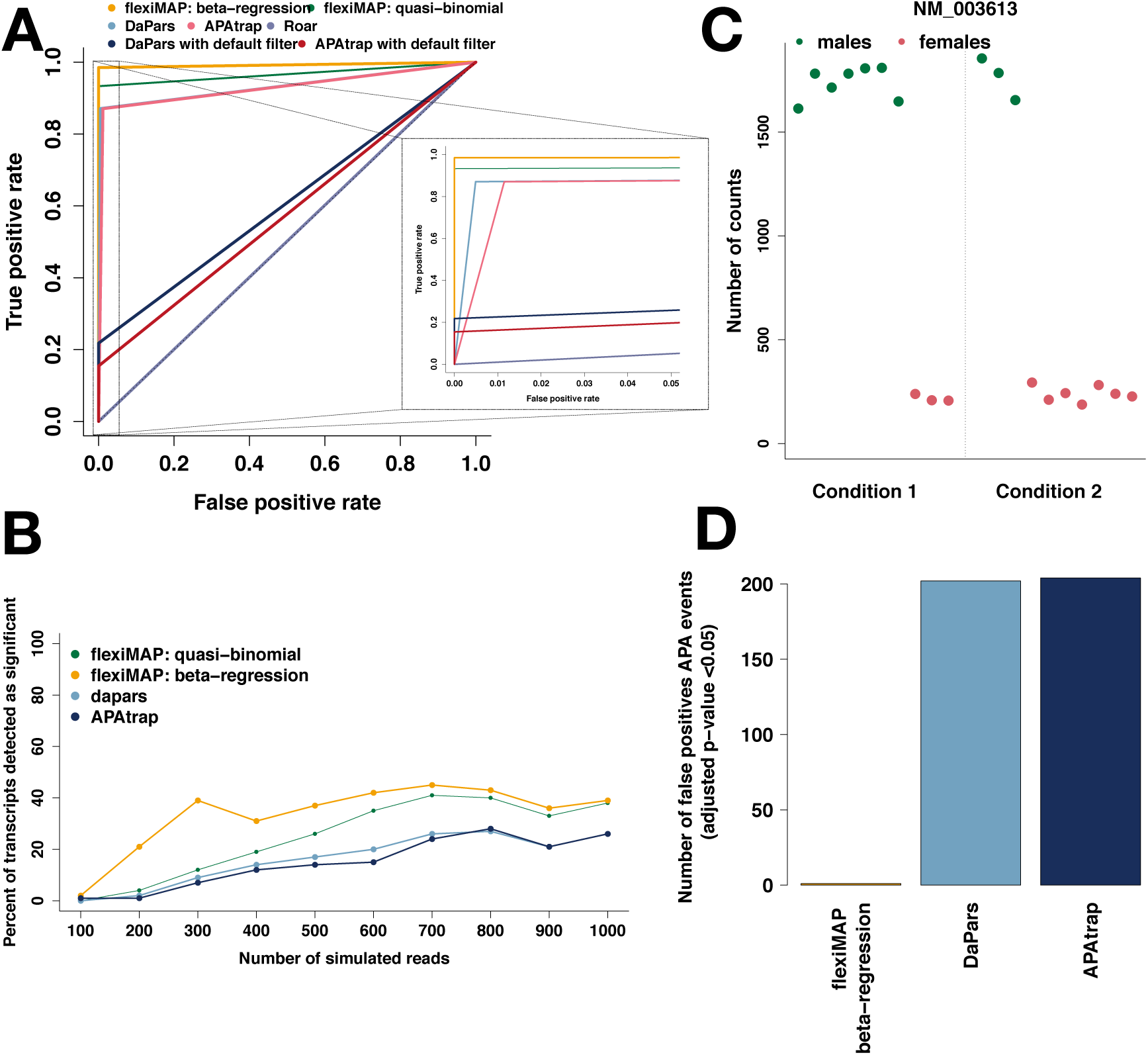
flexiMAP detects differential polyadenylation events with high specificity and outperforms in sensitivity DaPars and APAtrap at small fold changes. a) Receiver operating characteristic (ROC) curves representing the accuracy of detecting differential alternative polyadenylation events using flexiMAP, DaPars, APAtrap and Roar. Only transcripts where the polyadenylation site has been correctly predicted by DaPars and APAtrap are included in this plot. FlexiMAP clearly outperformed DaPars and APAtrap by perfect specificity and improved sensitivity. Although application of the PDUI post-hoc filter (DaPars) and PD filter (APAtrap) (in dark blue and dark red respectively) corrected the false positives problem of these methods, it did so at the cost of sensitivity. The Roar method essentially predicts every change to be significant so, unsurprisingly, its results do not differ from random guesses. b) Dependence of method sensitivity on the overall expression level of the transcript (only results for fold change=1.5 are shown). Transcripts with correctly predicted proximal sites whose APA events have been (correctly) identified as being significantly different between the two conditions show higher levels of expression across the whole 3’ UTR, whereas events that are missed originate in transcripts of lower overall expression. The beta-regression approach displays improved sensitivity, with the remaining methods showing the same overall trend but performing less well overall. c) Example from the imbalanced simulated dataset of a situation where a covariate of no interest (in this case, sex) affects the ratio of reads assigned to short and long isoforms. Male samples display much higher expression of the short region of transcript NM_003613 compared with female ones, regardless of the condition group samples belong to. In addition, the dataset is imbalanced, with more males present in condition 1 than condition 2. The mean expression for condition 1 is thus higher than the mean for condition 2, but the effect is due to the covariate sex, not the condition to which the samples belong to. d) DaPars and APAtrap report a large number of false positives for an imbalanced simulated dataset. In contrast, flexiMAP reports only one false positive in this case, highlighting its main advantage over alternative approaches.

We count RNA-seq reads falling in the “long” and “short” regions of the 3’ UTR (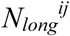 and 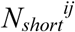 respectively), and define the ratio, *R*, for gene *i* in sample *j* as:

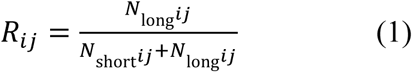

Reads falling in the long region can only originate from transcripts using the distal site, whereas reads falling in the short region may come from transcripts using either the distal or the proximal site. The ratio *R*_*ij*_ is the proportion of reads falling in the long region and is thus strictly contained in the interval (0,1). We note that the extreme value of zero is only encountered in the complete absence of a long isoform, whereas values greater than 0.5 would be observed only in cases where the long region is longer than the short region, or where strong 3’ biases in the read coverage are observed.

Our initial tests modeling APA events using logistic regression with quasi-binomial error distribution (within the Generalised Linear Model framework) showed that this approach was not sensitive enough for small numbers of samples or small fold changes. Different link functions did not improve the results. To allow more flexibility in modeling errors, we adopted instead a model where the response variable is assumed to be beta-distributed. This beta-regression model was implemented using the *betareg* package in *R* (Cribari-Neto and Zeileis, 2010). Results from the GLM approach with a quasi-binomial error family are included below for comparison.

Finally, our method incorporates two filtering steps to improve accuracy, employing TIN values (Wang *et al.*, 2012) to filter on transcript integrity and removing transcripts with too few reads mapping to the short region (see Supplemental Methods for details).

## Comparison to other methods using simulated data

We compared flexiMAP to three existing methods for APA analysis (DaPars (Xia *et al.*, 2014), Roar (Grassi *et al.*, 2016) and APAtrap (Ye *et al.*, 2018)) using simulated data we produced with the *polyester* R package (Frazee *et al.*, 2015) (see Supplementary Methods for details). Our method is specific (none of the transcripts with no APA events are predicted as having such events) and outperforms in sensitivity DaPars and APAtrap up to a fold change of 4 (Fig. 1A, Supp. Fig. 2). For larger fold changes, all methods appear to perform equally well. Surprisingly, the application of post-detection filters recommended by the developers of both DaPars and APAtrap appear to remove the majority of significant events across all fold changes, which renders questionable the usefulness of these filters (Fig. 1A). Finally, although Roar appears to be more sensitive than flexiMAP at small fold changes, in reality it predicts all transcripts as having APA events, regardless of whether they do or not, making its predictions unusable in practice (see also Supplementary Methods).

All methods, including flexiMAP, were sensitive to the expression level of the transcript tested for differential polyadenylation (Fig. 1B). APA events that were missed originated in transcripts of lower overall expression but the beta-regression approach displayed improved sensitivity.

Unlike methods that average across samples from the same condition, the performance of flexiMAP depends on the number of samples available in each group, as expected for a method that needs to model the variance within each group (see Supp. Fig. 3). However, flexiMAP is much more sensitive than the GLM-quasi-binomial method at small sample sizes (<6), often encountered in RNA-seq datasets.

The development of flexiMAP was primarily driven by the need to model multiple known covariates in APA analysis. Indeed, flexiMAP successfully discriminates between the effect of the condition of interest and that of an additional covariate in a simple simulated scenario of imbalanced datasets, where APA originates from the sex attribute of the samples rather than the condition of interest (Fig. 1C-D). Methods that cannot account for multiple covariates misinterpret the origin of this variation, resulting in increased false positive rates.

## Conclusion

We presented here flexiMAP, a beta-regression-based method for detecting alternative polyadenylation events in RNA-seq data, given a list of putative polyadenylation sites. Our method is both sensitive and specific, even when small numbers of samples are used, and has the distinct advantage of being able to model contributions from known covariates that would otherwise confound the results of APA analysis. flexiMAP compares favourably with existing alternatives in tests involving simulated datasets. Importantly, these tests have highlighted some hitherto overlooked caveats of existing methods.

## Acknowledgements / Funding

This work was supported by grants from the Birkbeck / Wellcome Trust Institutional Strategic Support Fund awarded to KJS and IN.

