## Supplementary Materials for "flexiMAP: A regression-based method for discovering differential alternative polyadenylation events in standard RNA-seq data"

#### Supplementary Methods

##### Description of the flexiMAP method

Given a catalogue of polyadenylation sites for a genome, the most downstream site per gene is selected as the end of the transcript and all other sites that are downstream from the coding region of a gene are potential alternative (proximal) polyadenylation sites. Each proximal site defines two regions in the 3' UTR (supplementary figure 1). The term "short" region is used here to refer to the part of the transcript starting from the start of the 3'UTR and ending at the proximal polyadenylation site and the term "long" region to refer to the part starting from the proximal site and ending at the most downstream polyadenylation site. It is important to emphasize that multiple proximal sites in one transcript are possible. All known sites belonging to one transcript are included in the flexiMAP analysis, but each proximal site is modeled separately.

Using the raw number of RNA-seq reads falling on each of these regions, the ratio $R$ is calculated for each transcript ($j$) in each sample ($i$) using:

$$R_{ij} = \frac{N_{\mathrm{long}ij}}{N_{\mathrm{short}ij} + N_{\mathrm{long}ij}}, \qquad (1)$$

where $N_{long}^{ij}$ and $N_{short}^{ij}$ are the number of reads falling in regions *long* and *short* respectively of transcript $i$ in sample $j$. It is worth pointing out here that reads falling in the long region can only originate from transcripts using the distal site, whereas reads falling in the short region may come from transcripts using either the distal or the proximal site. The ratio $R$ has the desired property of being constrained in the interval (0,1), where the extreme value 0 is only possible in the complete absence of the long isoform and values greater than 0.5 would generally only be observed if the long region was longer than the short region or if the dataset was affected by 3' or other biases in the distribution of reads (both of which could lead to a much larger number of reads covering the long region, compared with the short region).

Response variables representing proportions, like the ratio $R$ above, are commonly modelled using logistic regression. A link function (e.g. "logit") relates the mean of the response variable, $y$, to a linear predictor, $\eta$, and errors are usually assumed to be binomial. Count data in real experiments is often over-dispersed and in this case, the quasi-binomial family of errors can be used instead, thus doing away with the requirement of knowing in advance the relationship between mean and variance. In the context of APA, the main explanatory variable of interest is likely to be categorical (a "condition"), resulting in an ANOVA-like analysis of deviance.

In our own trials, modelling APA events using logistic regression with quasi-binomial error distribution (within the Generalised Linear Model framework in R) had poor sensitivity, when the number of samples was small or when small fold changes were involved. Application of different link functions did not improve the results. This suggested that perhaps the error distribution needed to be modeled more flexibly. Hence, we decided to adopt a model where the response variable is beta-distributed. The formula for the beta density, as parameterised by (Ferrari and Cribari-Neto, 2004) is given by the following equation:

$$f(y;\mu,\phi) = \frac{\Gamma(\phi)}{\Gamma(\mu\phi)\Gamma((1-\mu)\phi)} y^{\mu\phi-1}(1-y)^{(1-\mu)\phi-1} \quad , \qquad (2)$$

where $0 < \mu < 1$ is the mean of $y$ and $\phi$ is known as the precision parameter, with $\phi > 0$. $\Gamma$ denotes the gamma function. The variance of $y$ is given by $\mu(1-\mu)/(1+\phi)$, with precision parameter, $\phi$, allowing for wide range of shapes for the density. The beta distribution's density can have very different shapes depending of the value of the two parameters that determine the distribution, offering flexibility for modeling proportions. Furthermore, the interpretation of the results of a beta regression is similar to that of logistic regression with quasi-binomial errors. A beta-regression model was implemented here using the *betareg* package in R (Cribari-Neto and Zeileis, 2010). The package allows an extension to the original beta-regression model, where the precision parameter, $\phi$, is not assumed constant between observations but it is instead modeled in a similar fashion to the mean. However, we have not tested the use of a variable precision parameter as we believe the limited amount of data available would be problematic for modeling both parameters.

Our method currently relies on knowledge of the polyadenylation sites, usually obtainable from public databases. In a future version, the option of predicting sites from the density of RNA-seq reads could be added as a pre-processing step. However, this is not trivial. Even in simulated data, where read coverage of the 3'UTRs is perfect, DaPars predictions of proximal sites fall outside a 50-nucleotide window tolerance for around 40% of the sites (Supp. Fig. 4).

Our method incorporates two additional steps to improve accuracy. The first addresses the issue of distorted results in cases of severe RNA degradation levels. TIN values are pre-calculated as described in RSeQC (Wang *et al.*, 2012) to measure RNA integrity at transcript level and transcripts below a TIN threshold are removed. The second step addresses the problem of low expression by removing transcripts with fewer than a predefined threshold number of reads mapping to the "short" region.

A flow diagram of the flexiMAP method is shown in Supp. Fig. 5.

We note that adapting flexiMAP to be used for the analysis of data from long-read sequencing is relatively straightforward; relevant changes to the code are planned for future releases.

#### Details of simulated data

An "idealized" dataset of RNA-seq reads was created using the *polyester* R package (Frazee *et al.*, 2015). This simulated dataset is clean of technical biases and fold changes between isoforms are known, allowing testing of the sensitivity limits of the method in the absence of external factors. The simulation experiment comprises 20 samples, 10 in each of two conditions. Polyadenylation sites splitting each transcript into two isoforms (short and long) were obtained from the poly(A) site atlas (Gruber *et al.*, 2016) for 11000 human transcripts. Each isoform ("short" and "short + long") was simulated as a different transcript. The expression of the "shot + long" isoform was unchanged between conditions, whereas eleven different fold changes were applied between conditions for the "short" isoform in order to produce a range of different ratios, *R*. Hence, each fold change is represented by ~ 1000 transcripts in the dataset. Additionally, for each fold change category we assigned 100 different mean expression levels (from 100 to 1000) with the aim of sampling the effect of the expression level on the ability of the method to detect alternative polyadenylation events.

In order to assess the performance of flexiMAP in dealing with multiple covariates, an additional small dataset was simulated. The aim was to create a scenario where fold changes between two conditions are confounded by the presence of an additional factor. In the specific example set up, we created an imbalanced dataset where male and female-origin samples are present in unequal numbers in the control (7 males and 3 females) and condition (3 males and 7 females) groups. Although the group membership for the factor of interest (condition) plays no role in the choice of polyadenylation site of these transcripts, membership to male or female group does, confounding the outcome of methods that do not take into account additional covariates.

#### Quality control assessment of simulated data

A simulated dataset was used to assess the performance of flexiMAP. As differences in the length of the 3' UTRs across groups of transcripts could bias the results, we first checked the distribution profiles of the 3'UTR lengths across all the simulated groups (Supp. Fig. 4). Following filtering of transcripts with incorrectly predicted proximal polyadenylation sites by DaPars and APAtrap, on average, approximately 43% of transcripts were kept in each fold change group.

#### Mapping and counting reads in simulated data

Reads from the simulated dataset were mapped to the hg19 reference genome using HISAT2(Kim *et al.*, 2015). Uniquely mapped reads to non-overlapping genomic features were used as input to flexiMAP. Reads were counted using the *featureCounts* function from the Bioconductor package *Rsubread* (Liao *et al.*, 2014).

#### Assessing differential polyadenylation with DaPars

Once reads were mapped, the resulting SAM-formatted files were converted to the *bedgraph* format using Bedtools v2.17.0 (Quinlan and Hall, 2010) and these were used as input for the software DaPars-v.0.9 (Xia *et al.*, 2014). DaPars discovers statistically significant alternative polyadenylation events between two groups of samples. It predicts proximal sites using the drop in the number of reads near the site as a signal in a two-point model. In the case of simulated data, the proximal polyadenylation sites are already known and there is no need to predict them. Hence, in order to facilitate comparison between DaPars and other methods, we considered only cases where proximal polyadenylation sites were correctly predicted by DaPars (a correct prediction being defined here as a site that is located within 25 nucleotides in either direction of the known proximal site). Differences in the use of polyadenylation sites between treated and untreated samples were examined using the PDUI value, as defined by the software DaPars. The PDUI value is calculated for each gene passing the coverage thresholds of DaPars and is a measure of the preference of using the distal over the proximal poly(A) site (the DaPars software assumes and compares only two sites). If all reads are assigned to the distal site, the PDUI value is 1, and if all reads are assigned to the proximal site, 0. Mean PDUI values are calculated within each group and contrasted between groups to discover significant differential alternative polyadenylation events.

##### Assessing differential polyadenylation with APAtrap

A full analysis of the simulated dataset using APAtrap is estimated to take approximately 6 months (based on the estimator built into the software), so it was decided to apply this method only to the subset of transcripts with proximal polyadenylation sites correctly predicted by DaPars. The "Percentage Difference" (PD) is calculated by APAtrap to quantify the difference of APA site usage in a given gene between samples. A linear trend test based on the Pearson product moment correlation coefficient is employed to check the trend in proportions of APA usage and the false discovery rate is calculated using the Benjamini-Hochberg method.

##### Assessing differential polyadenylation with Roar

The method was applied to mapped simulated reads. The Roar ratio was calculated to quantify the difference of APA site usage in a given gene between samples. The Roar method assigns significance to the differences in APA site usage by carrying out pairwise comparison of samples between conditions using a Fisher exact test. Individual *p*-values from these pairwise comparisons are then multiplied by Roar to give an overall *p*-value. It is not at all obvious why this would be a valid approach and what would be the null hypothesis being tested by such a *p*-value. Given that this approach is almost guaranteed to lead to the majority of events (and likely all of them) being classed as significant, its practical value is also questionable.

#### Availability

The flexiMAP *R* package is available from:

https://github.com/kszkop/flexiMAP

Scripts and data to reproduce the analysis in this paper have been uploaded at the Zenodo repository (https://doi.org/10.5281/zenodo.3238619).

#### Supplementary Figure 1

**Definition of “short” and “long” region in the 3’ UTR of transcripts**

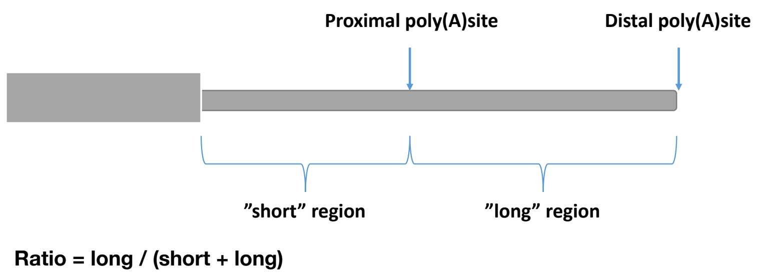


An alternative (proximal) poly(A) site in the 3’ untranslated region of a eukaryotic transcript splits this UTR into two parts, the “short” and the “long” regions. Isoforms terminating at the proximal site will only contain the short region whereas isoforms terminating at the distal site (often considered to be the canonical site) will contain both short and long regions. In this diagram the coding part of the exon is shown as a thick rectangle whereas the 3’ UTR is shown as the thinner rectangle.

#### Supplementary Figure 2

**flexiMAP detects differential polyadenylation events with high specificity and outperforms in sensitivity DaPars and APAtrap at small fold changes**

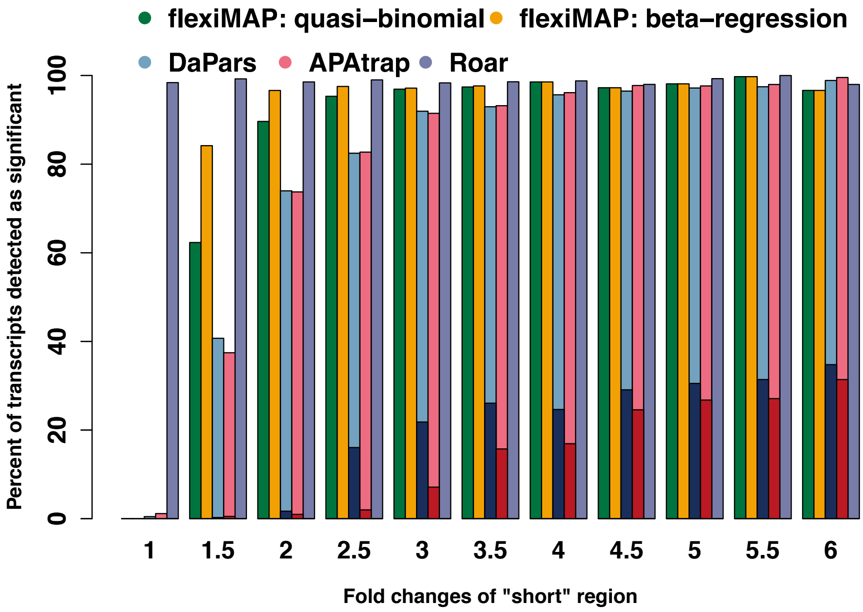


Percent of transcripts detected as significant (adjusted *p*-value <0.05) for every simulated fold change in the "short" region of the 3'UTR using flexiMAP, DaPars, APAtrap and Roar. Only transcripts where the polyadenylation site has been correctly predicted by DaPars and APAtrap are included in this plot. flexiMAP clearly outperformed Dapars and APAtrap for small fold changes. In addition, DaPars and APAtrap contain more false positives (see results for fold change=1). Although application of the recommended post-hoc filters (PDUI for DaPars and PD for APAtrap, in dark blue and dark red respectively) corrected the false positives problem, it did so at the cost of removing the majority of events from all remaining fold change categories. The Roar method essentially predicts every change to be significant so, unsurprisingly, it has very high sensitivity but its specificity is unacceptably low, rendering it unusable.

#### Supplementary Figure 3

**The flexiMAP beta-regression method is more sensitive than the GLM quasi-binomial approach for smaller numbers of samples per condition**

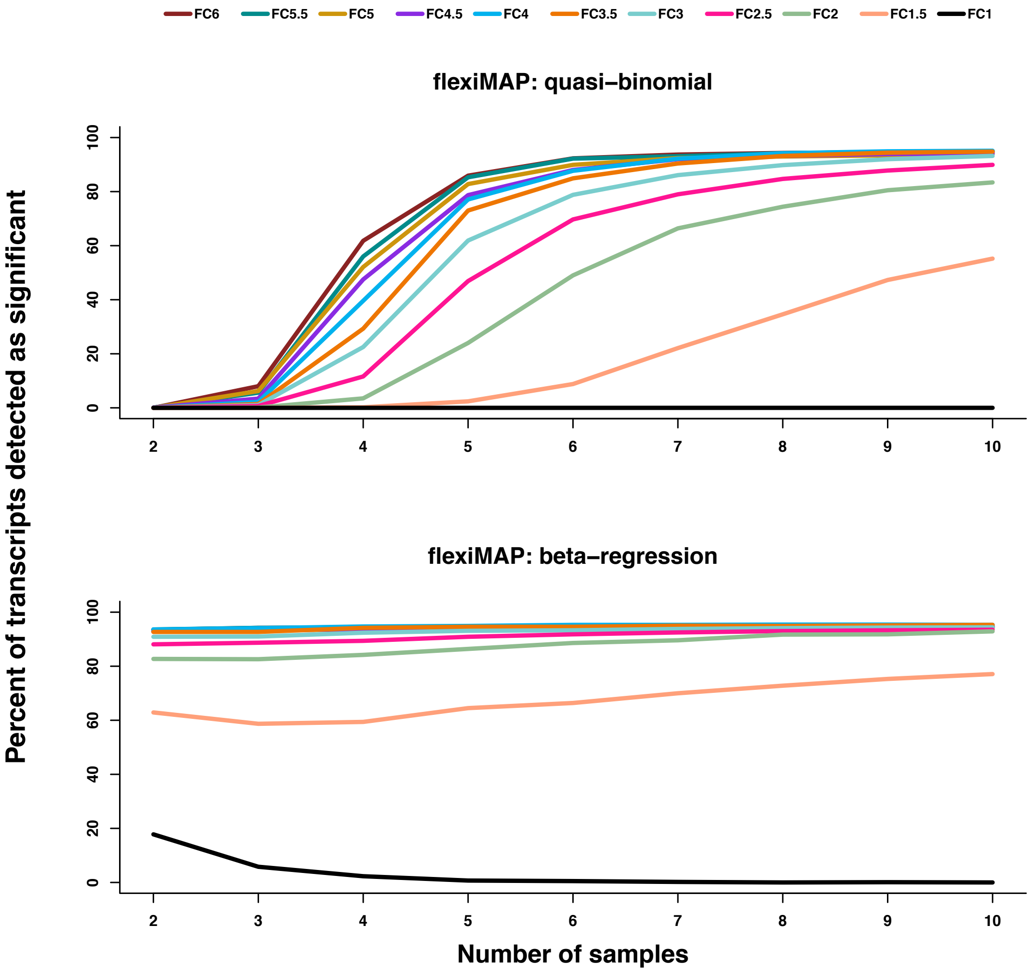


A GLM quasi-binomial approach detects hardly any APA events (even for highly expressed transcripts) when only a very small number of samples (<=3) per condition are available, indicating that this approach is not promising for the analysis of common RNA-seq datasets. Sensitivity is much improved by applying beta-regression but at the cost of a small fraction of false positives. DaPars and APAtrap results are similar; both are independent of the number of samples but require larger fold changes to display increased sensitivity.

### Supplementary Figure 4

#### Quality control assessment of simulated data

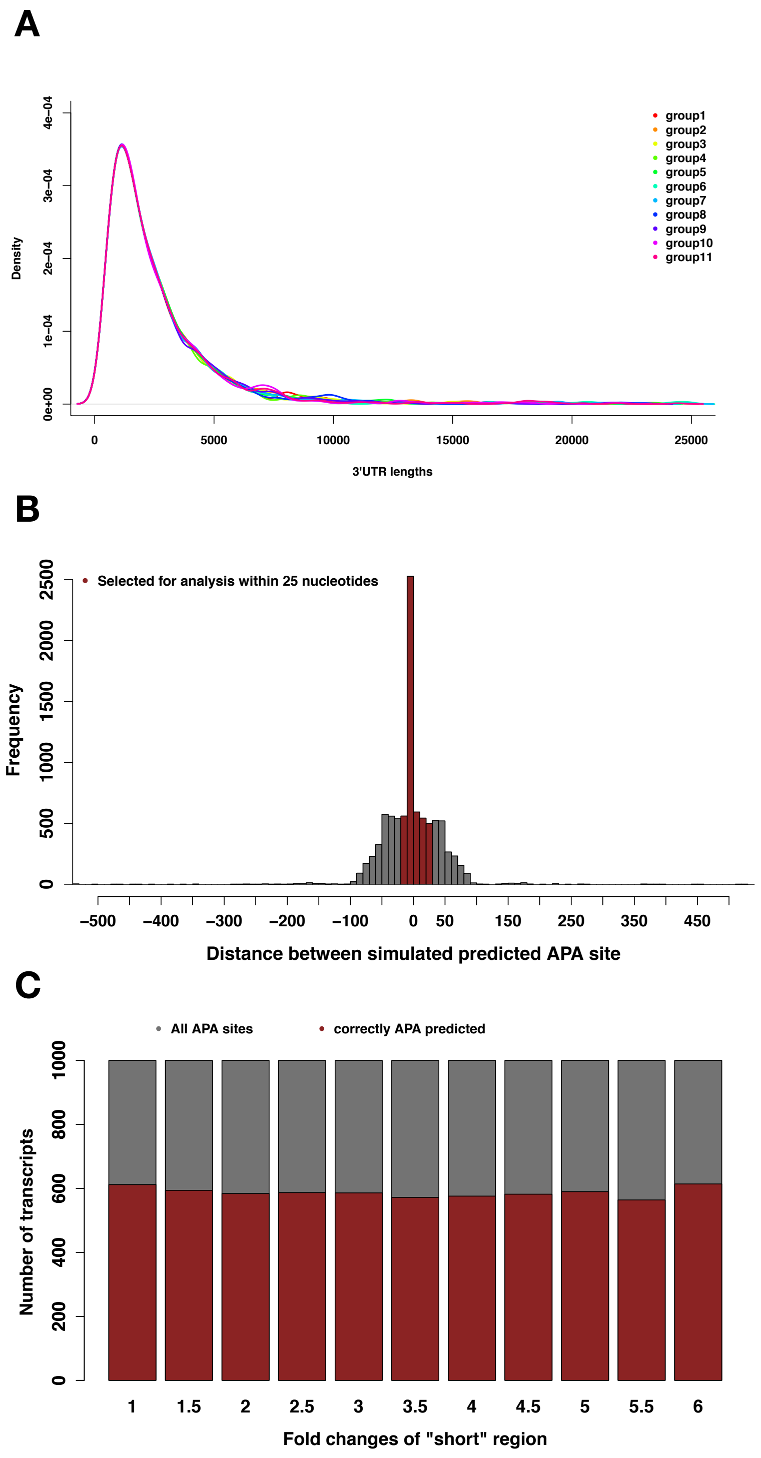

A) Density plots of the lengths of 3’UTR in each of the 11 groups in the simulated dataset indicates no length biases between the groups. Each group comprises 1000 transcripts and represents a different fold change.
B) Distribution of distances between simulated and predicted (by DaPars) proximal polyadenylation sites. Where the predicted site differed from the true site by up to 25 nucleotides in either direction, the prediction was considered correct and the corresponding transcript was kept in the simulation. These cases are highlighted in brown in the plot.
C) Transcripts with correctly predicted proximal polyadenylation sites are uniformly distributed across the fold change groups in the simulated dataset.

#### Supplementary Figure 5

##### Schematic diagram of flexiMAP workflow

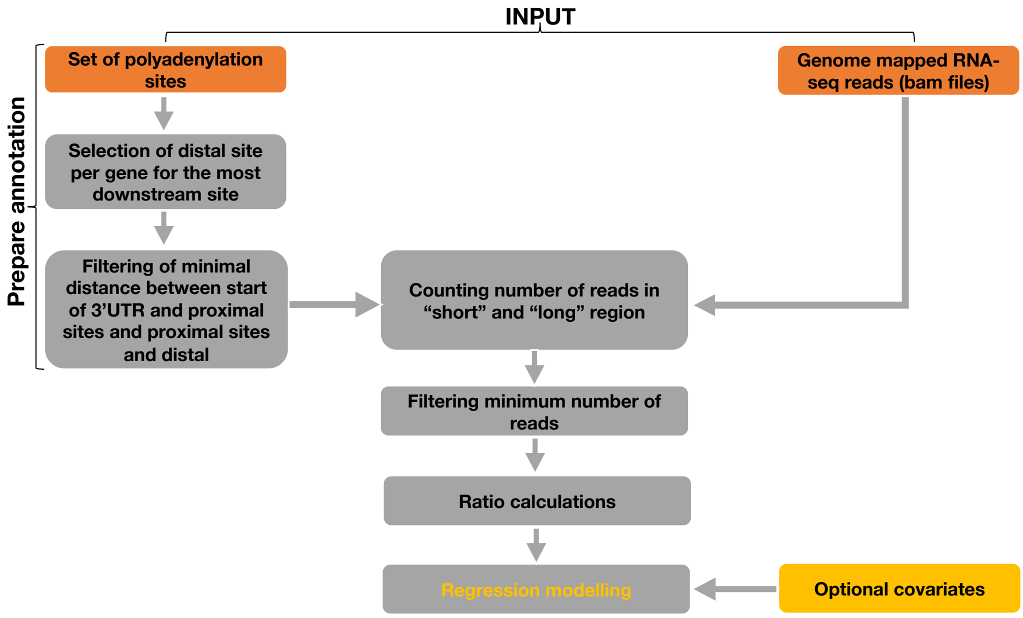
